# EPISeg: Automated segmentation of the spinal cord on echo planar images using open-access multi-center data

**DOI:** 10.1101/2025.01.07.631402

**Authors:** Rohan Banerjee, Merve Kaptan, Alexandra Tinnermann, Ali Khatibi, Alice Dabbagh, Christian Büchel, Christian W. Kündig, Christine S.W. Law, Dario Pfyffer, David J. Lythgoe, Dimitra Tsivaka, Dimitri Van De Ville, Falk Eippert, Fauziyya Muhammad, Gary H. Glover, Gergely David, Grace Haynes, Jan Haaker, Jonathan C. W. Brooks, Jürgen Finsterbusch, Katherine T. Martucci, Kimberly J. Hemmerling, Mahdi Mobarak-Abadi, Mark A. Hoggarth, Matthew A. Howard, Molly G. Bright, Nawal Kinany, Olivia S. Kowalczyk, Patrick Freund, Robert L. Barry, Sean Mackey, Shahabeddin Vahdat, Simon Schading, Stephen B. McMahon, Todd Parish, Véronique Marchand-Pauvert, Yufen Chen, Zachary A. Smith, Kenneth A. Weber, Benjamin De Leener, Julien Cohen-Adad

## Abstract

Functional magnetic resonance imaging (fMRI) of the spinal cord is relevant for studying sensation, movement, and autonomic function. Preprocessing of spinal cord fMRI data involves segmentation of the spinal cord on gradient-echo echo planar imaging (EPI) images. Current automated segmentation methods do not work well on these data, due to the low spatial resolution, susceptibility artifacts causing distortions and signal drop-out, ghosting, and motion-related artifacts. Consequently, this segmentation task demands a considerable amount of manual effort which takes time and is prone to user bias. In this work, we (i) gathered a multi-center dataset of spinal cord gradient-echo EPI with ground-truth segmentations and shared it on OpenNeuro https://openneuro.org/datasets/ds005143/versions/1.3.0, and (ii) developed a deep learning-based model, EPISeg, for the automatic segmentation of the spinal cord on gradient-echo EPI data. We observe a significant improvement in terms of segmentation quality compared to other available spinal cord segmentation models. Our model is resilient to different acquisition protocols as well as commonly observed artifacts in fMRI data. The training code is available at https://github.com/sct-pipeline/fmri-segmentation/, and the model has been integrated into the Spinal Cord Toolbox as a command-line tool.

## 1 Introduction

Functional magnetic resonance imaging (fMRI) allows non-invasive investigation of the central nervous system, of which, the spinal cord (SC) is a crucial part [Kinany et al., 2022, Powers et al., 2018, Haynes et al., 2023]. SC fMRI has the potential for tracking disease progression and assessing treatment effectiveness by providing novel insights into various neurological conditions such as multiple sclerosis, neuropathic pain, and SC injury [Chen et al., 2015, Landelle et al., 2023, Staud et al., 2021, Conrad et al., 2018]. Although fMRI was introduced in the 1990s, the application of fMRI in the SC has been lagging behind brain imaging studies due to the technical challenges owing to data acquisition and processing [Stroman et al., 2014, Cohen-Adad, 2017]. Despite these difficulties and thanks to recent technical developments, SC fMRI is rapidly taking off, with applications to study sensorimotor systems, as well as autonomic processes in humans and animals [Barry et al., 2014, Kinany et al., 2020, Wu et al., 2019, Tinnermann et al., 2022, K. A. Weber et al., 2020, Sengupta et al., 2024, Hemmerling et al., 2023, Vahdat et al., 2015, Khatibi et al., 2022, K. A. Weber et al., 2016, K. Weber et al., 2016, K. A. Weber et al., 2018, Kündig et al., 2024].

FMRI data are commonly acquired with a gradient-echo echo planar imaging (EPI) sequence, due to its short acquisition times (about 2s per volume) and its heavy T2* weighting, which produces increased sensitivity to the blood oxygenation level contrast [Stehling et al., 1991]. The most notable challenge arises from the substantial changes in the magnetic susceptibility of tissues surrounding the SC, which introduces image distortions and signal drop-outs in the images [Powers et al., 2018, Kinany et al., 2022, Stroman et al., 2014, Cohen-Adad, 2017]. Additionally, physiological noise, motion artifacts, and poor contrast between the SC and surrounding pulsatile cerebrospinal fluid lead to degradation in the quality of EPI images [Giove et al., 2004, Fratini et al., 2014].

The analysis of fMRI data typically involves statistical inferences at the group level, which necessitates the valid and precise normalization of data into a template space which aims to establish a one-to-one correspondence between the data from different individuals. The emergence of SC templates [De Leener et al., 2018, Fonov et al., 2014, Azzarito et al., 2020, Taso et al., 2014, Bosma and Stroman, 2014] and normalization procedures [De Leener et al., 2017, Stroman et al., 2008, Azzarito et al., 2020] make the group-level analysis accessible and reproducible across different SC fMRI datasets.

Currently, the gold standard for registering SC data to a template relies on the segmentation of the SC [Valosek and Cohen-Adad, 2024]. While there has been important development of automatic methods to segment the SC on MRI images [Blanc et al., 2023, De Leener et al., 2014, Gros et al., 2018, Bédard et al., 2023, Karthik et al., 2024, Horsfield et al., 2010, Hohenhaus et al., 2024, Tsagkas et al., 2023, Toufani et al., 2021, Lukas et al., 2021], these methods do not work well on gradient-echo EPI data due to aforementioned challenges. Moreover, the relatively low resolution of axial EPI scans (typically 1-1.5 mm in-plane, with 3-5mm thickness) makes the segmentation task even harder. An automated and robust segmentation method for the SC is therefore desirable to increase the reproducibility of SC fMRI studies [Hoggarth et al., 2022].

In this work, we introduce EPISeg, a deep learning (DL) based automatic SC segmentation method. Along with EPISeg, this paper also introduces open-access fMRI data of the SC from multiple sites (N=15). The dataset comes with ground-truth SC segmentations that were used to train the EPISeg model. We demonstrate the robustness of EPISeg amongst different sites, involving different scanners including data from healthy volunteers as well as patient groups, and against commonly observed artifacts such as partial volume effects, and ghosting. We compare our method with other state-of-the-art segmentation models like PropSeg [De Leener et al., 2014], DeepSeg [Gros et al., 2018] and Contrast-agnostic SC segmentation [Bédard et al., 2023], and show a significant improvement in segmentation performance using the Dice Score (DS) and Hausdorff Distance (HD). To our best knowledge, this is the first work regarding automatic segmentation of SC on fMRI data. The contributions of this paper are:

1. Creation of a new open-source SC gradient-echo EPI dataset (n=406 participants, 15 sites).
2. Introduction of an automated segmentation method for SC on fMRI data, thoroughly validated on diverse datasets spanning different populations and different regions of the SC including the lumbar and cervical SC.
3. Integration of this algorithm as a single-line command in SCT [De Leener et al., 2017], making it easily accessible to the scientific community.

## 2 Methods

In this section, we describe the data collection, manual labeling, model architecture, and model training using an active learning framework. An overview of the pipeline is shown in Figure 1.

**Figure 1:**
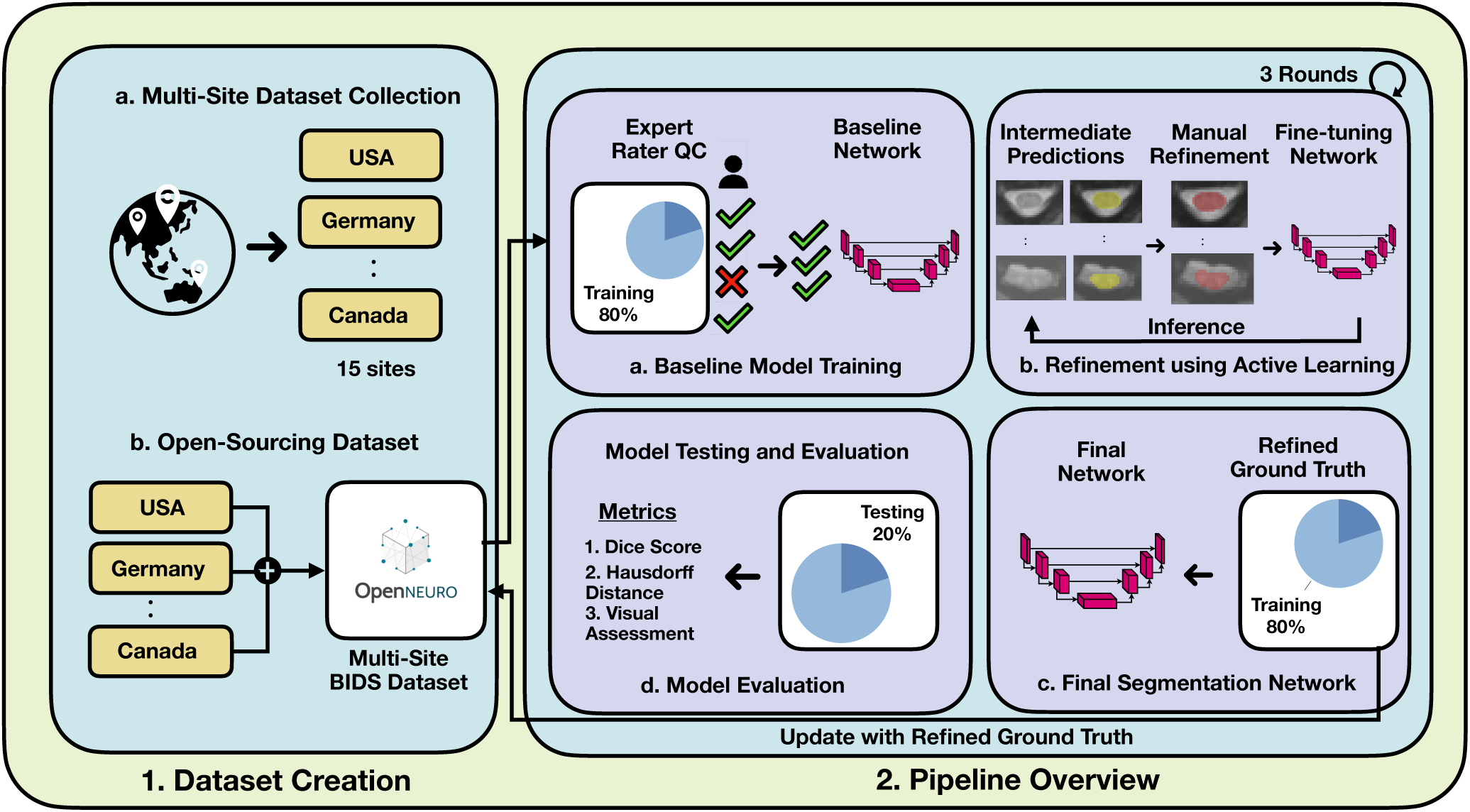
EPISeg overall pipeline. 1. Dataset creation and open-sourcing process. 1a. Initiation of dataset contribution call for EPISeg. 1b. The collected datasets are rearranged according to the brain imaging data structure (BIDS) convention and uploaded to OpenNeuro, forming a multi-site BIDS dataset. 2. Training and validation pipeline overview. 2.a. An expert selects reliable segmentations, which are used to train the baseline model. This model undergoes expert rater quality control (QC) to ensure high-quality training data. 2b. The baseline model generates initial predictions, which are then manually refined by experts. These refined predictions are used to fine-tune the baseline model. This process is repeated for three rounds, ensuring incremental improvement and robustness. 2c. Participants and refined ground truth from the refinement phase are used to train the final network. The refined ground truth is also updated in the open-source dataset. 2d. The final network is used for inference on test data, with metrics such as Dice Score (DS), Hausdorff Distance (HD), and visual assessments reported for comprehensive evaluation.

### 2.1 Data

We gathered gradient-echo EPI datasets from multiple sites (15 sites, 406 participants in total). All volunteers provided written informed consent following Institutional Review Board approval and the Declaration of Helsinki (see Table 1 for details of acquisitions for all sites). All participating sites were asked to share (i) the first 20 motion-corrected EPI volumes of the functional run (more volumes were unnecessary as the primary goal of these images is to perform segmentation, not fMRI analysis), (ii) the BIDS-compatible JSON file containing information about the acquisition, (iii) the mean image of the motion-corrected data, and (iv) the SC segmentation based on the mean motion-corrected image. Some sites manually created the segmentations from scratch, while other sites started off from a segmentation generated by automated methods such as PropSeg or DeepSeg, with manual correction. Out of 406 volunteers across all sites, the majority were healthy volunteers (n=359), 17 volunteers had degenerative cervical myelopathy, and 30 volunteers had fibromyalgia.

**Table 1:** Dataset details.

| Site name | Institution | Sample | Scanner and sequence |
| --- | --- | --- | --- |
| Leipzig_rest★<br>[Kaptan et al., 2022<br>Kaptan, Horn, et al., 2023] | Max Planck Institute for Human Cognitive and Brain Sciences, Leipzig, Germany | HV; N=45 (20F); Age: $27 \pm 3.8$ | 3T whole-body Siemens Prisma MRI System; 2D gradient-echo EPI; Number of slices: 24; in-plane resolution: $1.0 \times 1.0\text{mm}^2$; slice thickness: 5.0mm; FoV = $128 \times 128\text{mm}^2$; TR: 2.312s; TE: 40 ms; FA = $84^\circ$ |

Table 1: Dataset details (continued)
| Site name | Institution | Sample | Scanner and sequence |
| --- | --- | --- | --- |
| Geneva_rest1<br>[Kinany et al., 2019] | École Polytechnique Fédérale de Lausanne, Lausanne, Switzerland | HV; N=17 (11F); Age: $27 \pm 3.5$ | 3T whole-body Siemens Prisma MRI System; 2D gradient-echo EPI; number of slices: 27; FoV = $48 \times 144 \text{mm}^2$; in-plane resolution = $1 \times 1 \text{mm}^2$; slice thickness = 3mm; TR = 2.5s; TE = 34 ms; FA = $80^\circ$ |
| Geneva_rest2<br>[Kinany et al., 2020] | École Polytechnique Fédérale de Lausanne, Lausanne, Switzerland | HV; N=12 (6F); Age: $28.8 \pm 3.2$ | 3T whole-body Siemens Prisma MRI System; 2D gradient-echo EPI; number of slices: 27; FoV = $48 \times 144 \text{mm}^2$; in-plane resolution = $1 \times 1 \text{mm}^2$; slice thickness = 3mm; TR = 2.5s; TE = 34 ms; FA = $80^\circ$ |
| Geneva_rest3<br>[Kinany et al., 2022] | École Polytechnique Fédérale de Lausanne, Lausanne, Switzerland | HV; N=14 (7F); Age: $28.6 \pm 4.2$ | 3T whole-body Siemens Prisma MRI System; 2D gradient-echo EPI; number of slices: 31; FoV = $48 \times 144 \text{mm}^2$; in-plane resolution = $1 \times 1 \text{mm}^2$; slice thickness = 3mm; TR = 2.5s; TE = 34 ms; FA = $80^\circ$ |
| KCL_rest★<br>[Kowalczyk et al., 2024] | King's College London, London, UK | HV; N=23 (13F); Age: $23.9 \pm 3.8$ | 3T General Electric MR750 System; 2D gradient-echo EPI; number of slices: 38; FoV = 180mm; reconstruction matrix = $128 \times 128 \text{mm}^2$; in-plane resolution = $1.875 \times 1.875 \text{mm}^2$; slice thickness = 4mm (1mm gap); TR = 2.5s; TE = 30 ms; FA = $90^\circ$ |

Table 1: Dataset details (continued)
| Site name | Institution | Sample | Scanner and sequence |
| --- | --- | --- | --- |
| NW_motor<br>[Hemmerling<br>et al., 2023] | Northwestern Uni-<br>versity, Chicago,<br>Illinois, USA | HV; N=28<br>(18F); Age:<br>26±4.7 | 3T whole-body Siemens Prisma MRI<br>System; 2D gradient-echo EPI; num-<br>ber of slices: 25; FoV = 128 ×<br>44mm<sup>2</sup>; in-plane resolution = 1 ×<br>1mm<sup>2</sup>; slice thickness = 3mm; TR =<br>2s; TE = 30ms; FA = 90° |
| NW_motor-Weber<br>[K. A. We-<br>ber et al.,<br>2016] | Northwestern Uni-<br>versity, Chicago,<br>Illinois, USA | HV; N=11<br>(6F); Age:<br>27.7±1.9 | 3T whole-body Siemens Prisma MRI<br>System; 2D gradient-echo EPI; num-<br>ber of slices: 31; FoV = 128 ×<br>44mm<sup>2</sup>; in-plane resolution = 1 ×<br>1mm<sup>2</sup>, slice thickness = 3mm, TR =<br>2.5s, TE = 30ms, FA = 80° |
| NW_thermal<br>[K. Weber<br>et al., 2016] | Northwestern Uni-<br>versity, Chicago,<br>Illinois, USA | HV; N=12<br>(3F); Age:<br>28.8±2.5 | 3T whole-body Siemens Prisma MRI<br>System; 2D gradient-echo EPI; num-<br>ber of slices: 31; FoV = 128 ×<br>128mm<sup>2</sup>; in-plane resolution = 1 ×<br>1mm<sup>2</sup>, slice thickness = 3mm, TR =<br>2.5s, TE = 30ms, FA = 80° |
| NW_tactile<br>[K. A. We-<br>ber et al.,<br>2020] | Northwestern Uni-<br>versity, Chicago,<br>Illinois, USA | HV; N=12<br>(3F); Age:<br>28.8±2.5<br>Degenerative<br>cervical<br>myelopathy<br>N = 2; Age:<br>33.5 ± 2.1 | 3T whole-body Siemens Prisma MRI<br>System; 2D gradient-echo EPI; num-<br>ber of slices: 25; FoV = 128 ×<br>44mm<sup>2</sup>; in-plane resolution = 1 ×<br>1mm<sup>2</sup>; slice thickness = 3mm; TR =<br>2s; TE = 30ms; FA = 80° |

Table 1: Dataset details (continued)
| Site name | Institution | Sample | Scanner and sequence |
| --- | --- | --- | --- |
| Zurich_Cerv<br>_Bilatmotor | Balgrist Univer-<br>sity Hospital,<br>University of<br>Zurich, Zurich,<br>Switzerland | HV; N=18<br>(3F); Age:<br>28.8±2.5 | 3T whole-body Siemens Prisma MRI<br>System; 2D gradient-echo EPI; num-<br>ber of slices: 18; FoV = 128 ×<br>128mm<sup>2</sup>; in-plane resolution = 1 ×<br>1mm<sup>2</sup>; slice thickness = 5mm; TR =<br>3.26s; TE = 31ms; FA = 87° |
| Stanford_rest<br>[Kaptan,<br>Law, et al.,<br>2023] | Stanford Univer-<br>sity, Palo Alto,<br>California, USA | HV; N=29<br>(F); Age:<br>25.6±5.5 | GE 3T Discovery 750; 2D gradient-echo EPI; number of slices: 43; FoV<br>= 80mm; matrix = 64 × 64mm<sup>2</sup>; in-plane resolution = 1.25 × 1.25mm<sup>2</sup>;<br>slice thickness = 5mm; TR = 2.5s;<br>TE = 30ms; FA = 80° |
| Stanford_<br>restmartucci<br>[Martucci<br>et al., 2021,<br>Martucci<br>et al., 2019] | Stanford Univer-<br>sity, Palo Alto,<br>California, USA | HV; N=15<br>(F); Age:<br>49±10.8<br>Fibromyalgia<br>N=30<br>(F); Age:<br>50.2±8.8 | 3T General Electric MR750 Sys-<br>tem; number of slices: 14; FoV =<br>160mm; reconstruction matrix = 128<br>× 128mm<sup>2</sup>; in-plane resolution =<br>1.25 × 1.25mm<sup>2</sup>; slice thickness =<br>4mm<sup>2</sup>; TR = 2.5s; TE = 30ms; FA<br>= 75° |
| UNF_MSL<br>[Khatibi<br>et al., 2022,<br>Mobarak-Abadi et al.,<br>2024] | Centre de<br>recherche de<br>l'Institut Universi-<br>taire de Gériatrie<br>de Montréal,<br>Montréal, QC,<br>Canada | HV; N=27<br>(14F); Age:<br>24.9±3 | 3T whole-body Siemens TIM Trio<br>MRI System; 2D gradient-echo; num-<br>ber of slices: 40-43 (8-10 SC); FoV =<br>132 × 132mm<sup>2</sup>; in-plane resolution =<br>1.2 × 1.2mm<sup>2</sup>; slice thickness = 5mm<br>(9mm gap); TR = 3.14s; TE = 33ms;<br>FA = 90° |

Table 1: Dataset details (continued)
| Site name | Institution | Sample | Scanner and sequence |
| --- | --- | --- | --- |
| OUHSC_stim | University of Oklahoma Health Sciences Center, OK, USA | HV; N=13(9F); Age: 50.3±6.6<br>Cervical Spondylotic Myelopathy<br>N=15(10F); Age: 57±9.9 | 3T whole-body Siemens Prisma MRI System; 2D gradient-echo EPI; number of slices: 10; FoV = 128 × 128mm<sup>2</sup>; in-plane resolution = 1 × 1mm<sup>2</sup>; slice thickness = 5mm; TR = 0.86s; TE = 30ms; FA = 50° |
| Hamburg_pain◆<br>[Tinnermann et al., 2021] | Department of Systems Neuroscience, University Medical Center Hamburg-Eppendorf, Hamburg, Germany | HV; N=30 (M); Age: 23.9±3.8 | 3T whole-body Siemens Prisma MRI System; 2D gradient-echo EPI; number of slices: 10; FoV = 128 × 128mm<sup>2</sup>; in-plane resolution = 1 × 1mm<sup>2</sup>; slice thickness = 5mm; TR = 0.86s; TE = 30ms; FA = 50° |
| Zurich_Lumb_rest<br>[Kündig et al., 2024] | Balgrist University Hospital, University of Zurich, Zurich, Switzerland | HV; N=13 (4F); Age: 28.7±3.5 | 3T whole-body Siemens Prisma MRI System; 2D gradient-echo EPI; number of slices: 15; FoV = 144 × 48mm<sup>2</sup>; in-plane resolution = 1 × 1mm<sup>2</sup>; slice thickness = 5mm; TR = 1.29s; TE = 35ms; FA = 74° |
| Leipzig_pain◆<br>[Dabbagh et al., 2024] | Max Planck Institute for Human Cognitive and Brain Sciences, Leipzig, Germany | HV; N=40 (20F); Age: 27.3±4.6 | 3T whole-body Siemens Prisma MRI System; 2D gradient-echo EPI; number of slices: 16; in-plane resolution: 1.0 × 1.0mm<sup>2</sup>; slice thickness: 5.0mm; FoV: 128 × 128mm<sup>2</sup>; TR: 1.8s; TE: 40ms; FA = 70° |

\*HV = Healthy Volunteers; FoV = Field of View; TR = Repetition Time; TE = Echo Time; FA = Flip Angle; ★ = Semi-automatic segmentation; ◆ = Automatic segmentation

The dataset was put together into one multi-site dataset and organized according to the BIDS convention [Gorgolewski et al., 2016]. The segmentations produced at each site were reviewed by three raters overseeing the entire study (RB, MK, JCA) using the sct qc quality control (QC) tool [Valosek and Cohen-Adad, 2024]. Segmentations were subsequently updated using EPISeg wherever we found erroneous segmentations. The dataset was uploaded on OpenNeuro (https://openneuro.org/datasets/ds005143/versions/1.3.0), except for one site due to the data sharing policy of their respective ethics committee.

### 2.2 Data preprocessing

Our proposed model is designed to segment the SC on fMRI images. The input consists of a mean motion-corrected volume representing the fMRI scan. This mean volume condenses the motion-corrected time-series data into a single three-dimensional scan by averaging over all time points. Each input volume is matched with a ground truth binary segmentation that outlines the spinal cord’s boundaries.

### 2.3 Criteria for selecting training data

When performing multi-step active learning training, the first training batch necessitates reliable ground truth segmentation. In this case, we aimed to identify data subsets, *D_g_*, that had accurate ground truth SC segmentations. These data subsets were in general of high quality, notably thanks to the z-shimming method [Finsterbusch et al., 2012] used in some of the datasets like the Leipzig pain, Leipzig rest. It is important to note that all selection processes prioritized accurate segmentations over image quality (which is inherently related to segmentation quality), ensuring that the model was trained on the most precise delineations of the SC.

### 2.4 Model training protocol

Our training protocol involves a three-step model training procedure employing a human-in-the-loop active learning to iteratively remove errors and improve ground truth accuracy and then training a final segmentation model which is deployed in SCT. Our reasoning for using active learning in this setting was to develop a model resilient to the various types of image artifacts it encounters over the different rounds of training.

In the first step, as shown in Figure 2 we trained a baseline model *M_b_* using the dataset *D_g_* which had 96 participants. The baseline model is a 3D U-Net [Çiçek et al., 2016], trained using the nnUNetv2 framework [Isensee et al., 2020], which self-configures parameters for optimal performance. We chose nnUNetv2 to perform this task because it has shown state-of-the-art performance in a range of medical anatomies and imaging modalities [Isensee et al., 2020, Isensee et al., 2024]. For training *M_b_*, we used the default parameters configured by nnUNetv2 for our dataset. The training details include a batch size of 5, learning rate of 0.01 with a polynomial learning rate scheduler and a stochastic gradient descent optimizer for a total of 1,000 epochs. The model was trained using the combination of Dice loss and cross-entropy loss.

**Figure 2:**
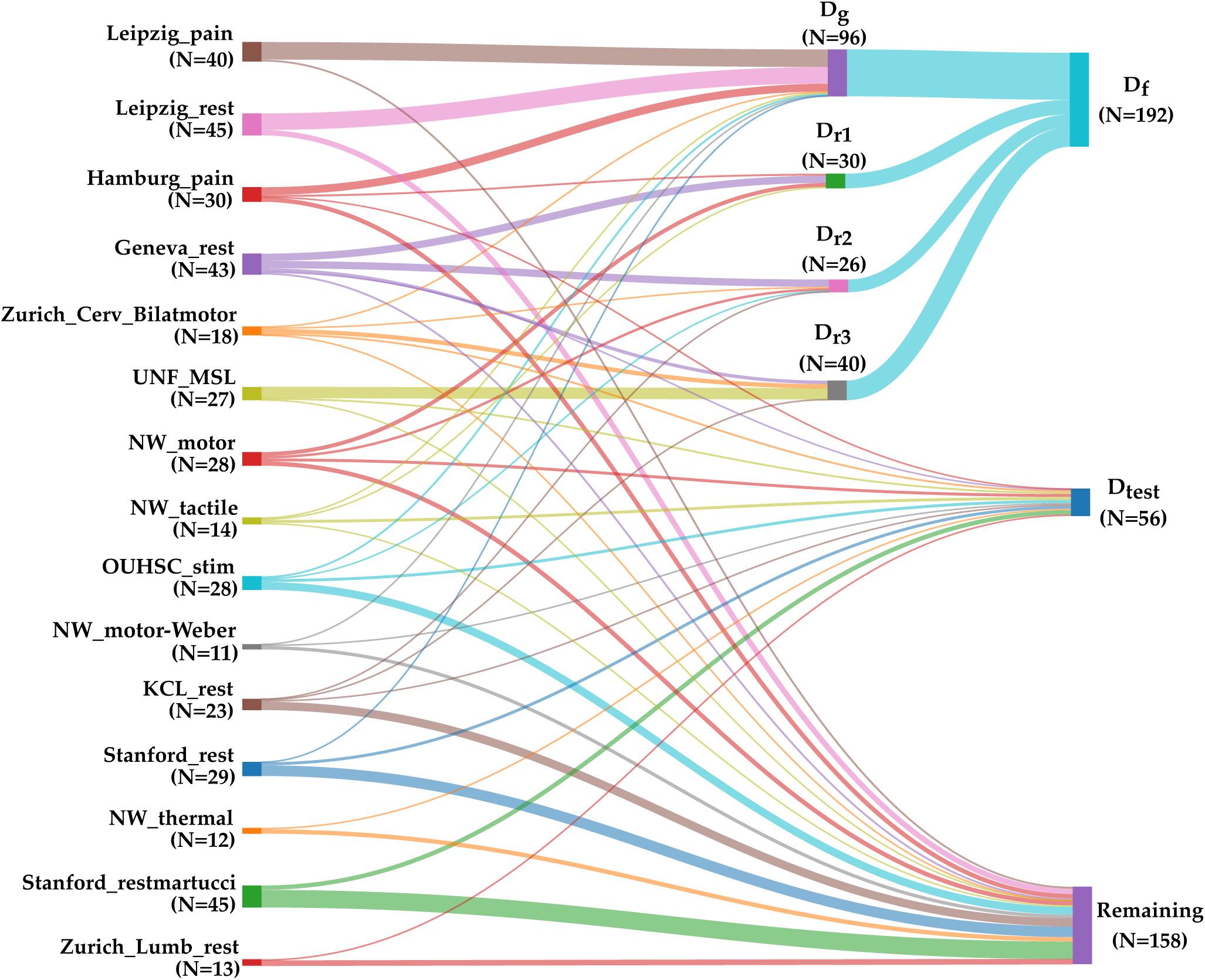
Dataset split. The sankey chart shows the different dataset splits created during the training process. The split *D_g_* is for the baseline model, *D_r_*_1_ is for the first round of active learning training, *D_r_*_2_ is for the second round of active learning training, *D_r_*_3_ is for the third round of active learning training, *D_f_* is the aggregated set of *D_g_*, *D_r_*_1_, *D_r_*_2_. *D_r_*_3_ and *D_test_* is the held-out test set to validate the EPISeg performance. The “remaining” set contains images with artifacts, erroneous segmentations.

The second step consisted of iterative refinement of training ground truth and model using active learning. Once the baseline model, *M_b_* was trained, we used it to generate segmentations on the remaining dataset. For the first round of active learning, a subset of these segmentations, *D_r_*_1_, consisting of 30 images were chosen for quality control assessment and manual corrections, if necessary. This subset *D_r_*_1_ was then used to fine-tune *M_b_*, resulting in an updated model in round 1 of training, *M_r_*_1_. To further improve the model’s capability, two more rounds of active learning with the same procedure with variable dataset sizes were conducted which resulted in the creation of datasets *D_r_*_2_, *D_r_*_3_ and corresponding models *M_r_*_2_ and *M_r_*_3_ respectively. These training rounds used the same default parameters used to train *M_b_*.

For the third and final step of the training protocol, all the datasets *D_g_*, *D_r_*_1_, *D_r_*_2_ and *D_r_*_3_ were aggregated into one final dataset, *D_f_* which had 192 3D images to train one model, *M_f_*. Our reasoning behind training the *M_f_* was to overcome the signs of forgetting that *M_r_*_3_ was exhibiting. Forgetting is a common phenomenon that is observed during active learning i.e. when networks are trained in a sequential fashion, the new learnings of the network tend to interfere with the old learning of the network [McCloskey and Cohen, 1989]. We discuss further about this observation in the “Effect of active learning on EPISeg” results section. Following a similar training procedure from the first step, a new set of nnUNetv2 parameters were configured for *D_f_*, which included a batch size of 15 with a polynomial learning rate scheduler, trained for 1,000 epochs. Along with this, we use the latest nnUNetv2 ResEnc [Isensee et al., 2024] variant of the UNet with residual connections in the encoder since the authors mention that using residual connections along with the nnUNetv2 framework is a recipe for better segmentation performance.

### 2.5 Evaluation protocol

To evaluate our model performance, we created a multi-site held-out test set, *D_test_*. The whole dataset (containing all subjects from all sites) was split in a 80:20 ratio where participants from the 80% of the dataset were used for training and 20% of the dataset were held out for testing. The *D_test_* was not included in the training or validation sets. To have a fair representation of all the sites in *D_test_*, we randomly chose 20% of the participants from each site and aggregated it into one dataset which in total had 56 participants. We evaluate our model’s performance on the *D_test_* and compare it with the SC segmentation models PropSeg, DeepSeg and Contrast-agnostic SC segmentation model. We discuss these results in the following sections.

We use Dice Score (DS) and Hausdorff Distance (HD) as validation metrics. For HD, we use the 95th percentile HD (HD95) since it reduces the impact of outlier errors, providing a more robust and representative measure of segmentation accuracy. The DS quantifies the spatial agreement between the predicted and ground truth segmentations, with higher scores indicating a closer alignment. On the other hand, the HD measures the maximum separation between the predicted SC segmentation and the nearest point in the ground truth segmentation, providing insights into the model’s ability to capture the geometric correspondence between predicted and true SC boundaries. We utilize the implementations provided by the MetricsReloaded framework [Maier-Hein et al., 2022] for our metrics. Additionally, we verified the model’s performance to ensure that model was producing segmentations as expected [discussed in section 3.4]

## 3 Results

### 3.1 Automatic fMRI SC segmentation

Table 2 compares the performance of the EPISeg models trained on various datasets - *M_r_*_3_ trained on the final round of the active learning data subset *D_r_*_3_; *M_f_* trained on the aggregated set *D_f_* from all rounds of active learning rounds; *M_ResEnc_* is the model with ResNet connections trained on *D_f_*. The best performance (in terms of DS) is seen with the *M_f_* dataset, indicating the advantage of having a single model with reliable segmentations rather than having several rounds of fine-tuning.

**Table 2:** Performance of the proposed EPISeg on *D_test_*. *M_r_*_3_ refers to the model we obtained after the 3*^rd^* round of active learning training, *M_f_* refers to the model trained on *D_f_* which is the final aggregated data from all the 4 rounds of active learning training and *M_ResEnc_*refers to latest ResNet based nnUNet model trained on *D_f_*.

| Model | DS ± S.D. | HD95 ± S.D. (in mm) |
| --- | --- | --- |
| EPISeg ($M_{r3}$) | 0.82 ± 0.06 | 1.73 ± 1.03 |
| EPISeg ($M_f$) | 0.88 ± 0.05 | 1.41 ± 0.86 |
| EPISeg ($M_{ResEnc}$) | 0.87 ± 0.05 | 1.28 ± 0.73 |

Even though we see *M_f_* performs a little better than *M_ResEnc_* in terms of DS, *M_ResEnc_* has a better HD95 score. When we investigated this qualitatively, we saw that *M_ResEnc_*gave more complete segmentations, for example, it was able to segment the first and last slice more consistently and was more robust to slices with artifacts. We therefore used the *M_ResEnc_* as our final EPISeg model.

### 3.2 Effect of active learning on EPISeg

We relied heavily on human-in-the-loop active learning for training EPISeg. Our selection of *D_g_* was based on the quality of the segmentations rather than the image quality. However, since it was easier to achieve better segmentations on higher quality images, they were predominantly used in training *M_b_*. More and more lower quality images were introduced in the later rounds of active learning. As we can observe in Figure 3, there is a noticeable dip in the performance of *M_r_*_3_. When we investigated this phenomenon, we observed that, while the model’s performance improved for lower quality images, it degraded the performance for high quality images similar to the ones used in training *M_b_*. We refer to this as forgetting and therefore decided to train *M_f_* to overcome this drawback of EPISeg.

**Figure 3:**
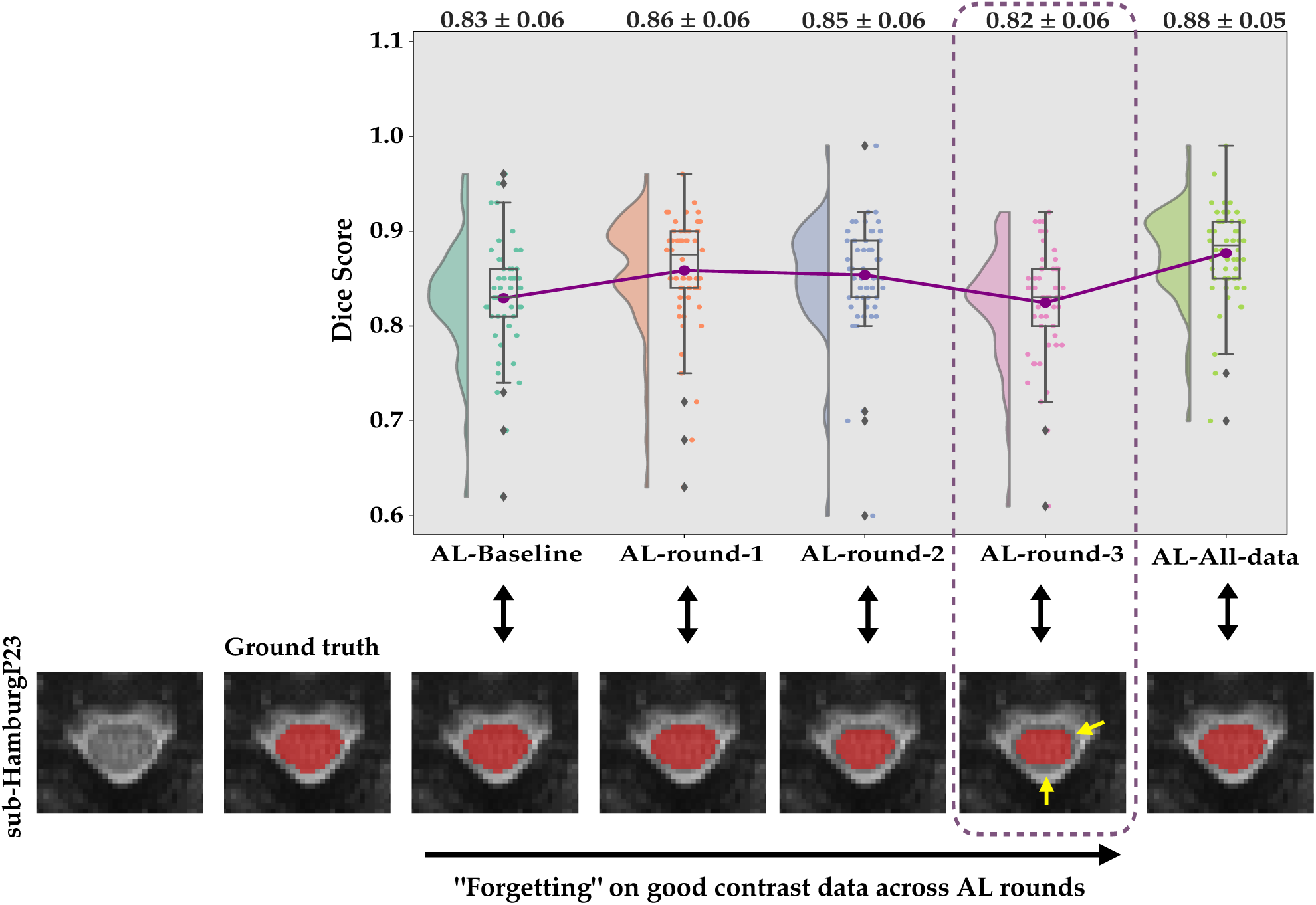
Effects of Active Learning. The raincloud plot shows the DS distribution of different models trained during the active learning training rounds. The images below show the axial slice from a subject and the red overlays show the model prediction of the SC. The dotted line and the yellow arrows show the performance degradation of the model over the training rounds.

### 3.3 Comparison with other methods

We compared EPISeg with other methods, namely, PropSeg, DeepSeg and Contrast-agnostic (from SCT v6.3: https://github.com/spinalcordtoolbox/spinalcordtoolbox/releases/tag/6.3). PropSeg does not use DL for SC segmentation and has been tested only on T1, T2 and T2* contrasts, DeepSeg uses DL but no EPI data were used during the training. Similarly, Contrast-agnostic did not use gradient-echo EPI data during training. However, both DeepSeg and Contrast-agnostic can segment EPI data. Our motive to compare our results with these methods was the fact that they are considered state-of-the-art, fully automatic 3D SC segmentation methods.

In Table 3, the various model performances on *D_test_* are presented. The proposed model, EPISeg, exhibits superior performance in comparison to the alternative baselines. Both PropSeg and DeepSeg depict subpar performance when dealing with EPI data. Numerous instances exist where these models struggle to accurately segment the SC. PropSeg, for example, faces difficulties in identifying the SC edges in many scenarios, resulting in segmentation failure. Conversely, Deepseg initially locates the center of mass before proceeding with SC segmentation, a task it struggles with in EPI SC segmentation. The Contrast-agnostic model demonstrates satisfactory but lower performance on our dataset due to its exposure to diverse SC contrasts during the training process.

**Table 3:** Comparison of SC segmentation methods on *D_test_*.

| Model | DS ± S.D. | HD95 ± S.D. (in mm) |
| --- | --- | --- |
| Propseg | $0.56 \pm 0.22$ | $4.70 \pm 3.81$ |
| Deepseg | $0.77 \pm 0.17$ | $2.08 \pm 1.38$ |
| Contrast-agnostic | $0.83 \pm 0.08$ | $1.76 \pm 1.33$ |
| EPISeg ($M_{r3}$) | $0.82 \pm 0.06$ | $1.73 \pm 1.03$ |
| EPISeg ($M_f$) | **$0.88 \pm 0.05$** | $1.41 \pm 0.86$ |
| EPISeg ($M_{ResEnc}$) | $0.87 \pm 0.05$ | **$1.28 \pm 0.73$** |

Figure 4 presents raincloud plots comparing the performance of different methods on *D_test_*. The half-violin plots illustrate the distribution of DS, while individual points represent the performance of each data point in *D_test_*. EPISeg’s DS are tightly clustered around a single point, which coincides with both the mean and median of the scores. This concentration indicates that EPISeg delivers consistent performance without outliers. When it comes to comparison of HD95 in Figure 5, we observe a similar pattern as in Figure 4. To highlight the strong performance of EPISeg, we see a point where the HD95 value is 0.0mm which, means that 95% or more surface points had 0.0mm error, indicating a perfect segmentation performance.

**Figure 4:**
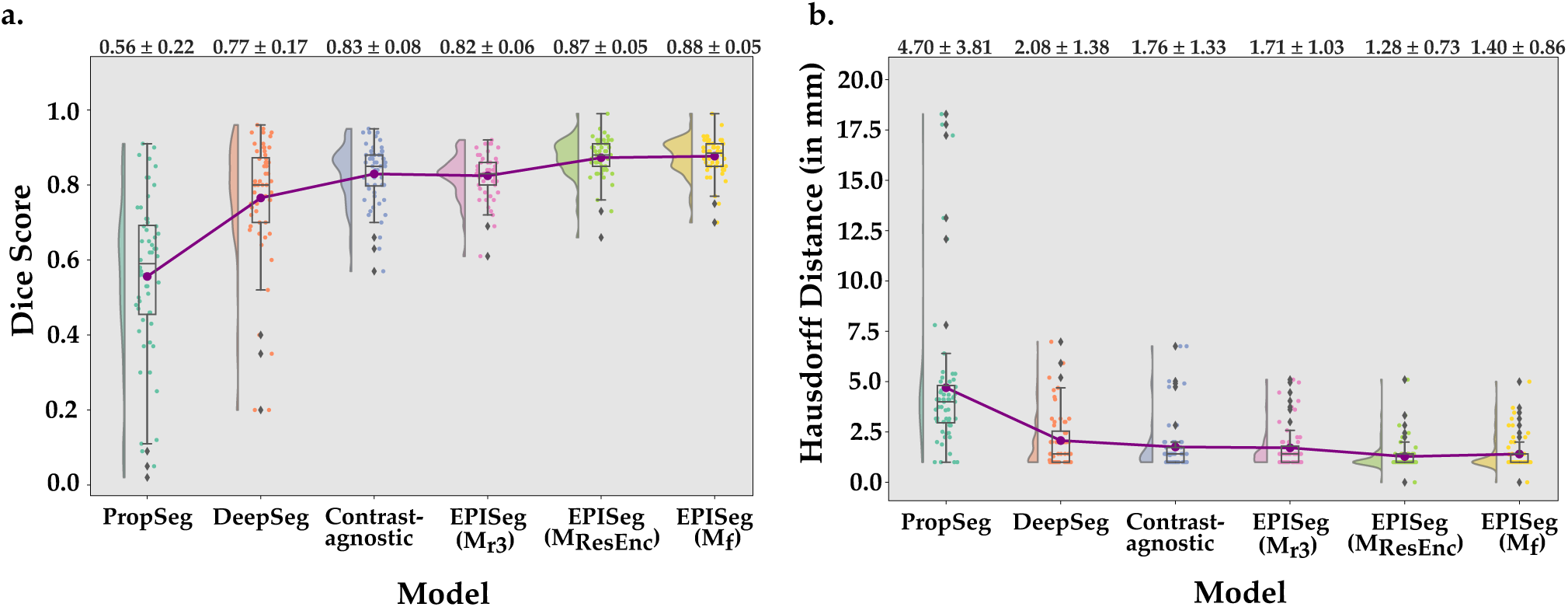
Raincloud plots from method comparison. 4a. The DS were computed between the ground truth and predictions. The raincloud plots show the DS distributions of the different models. The red dots and the line connect the mean values of the DS of the respective models. The mean DS along with the standard deviation is printed on the top of the plot. 4b. In a similar way to 4a., it shows the HD95 for the different models.

**Figure 5:**
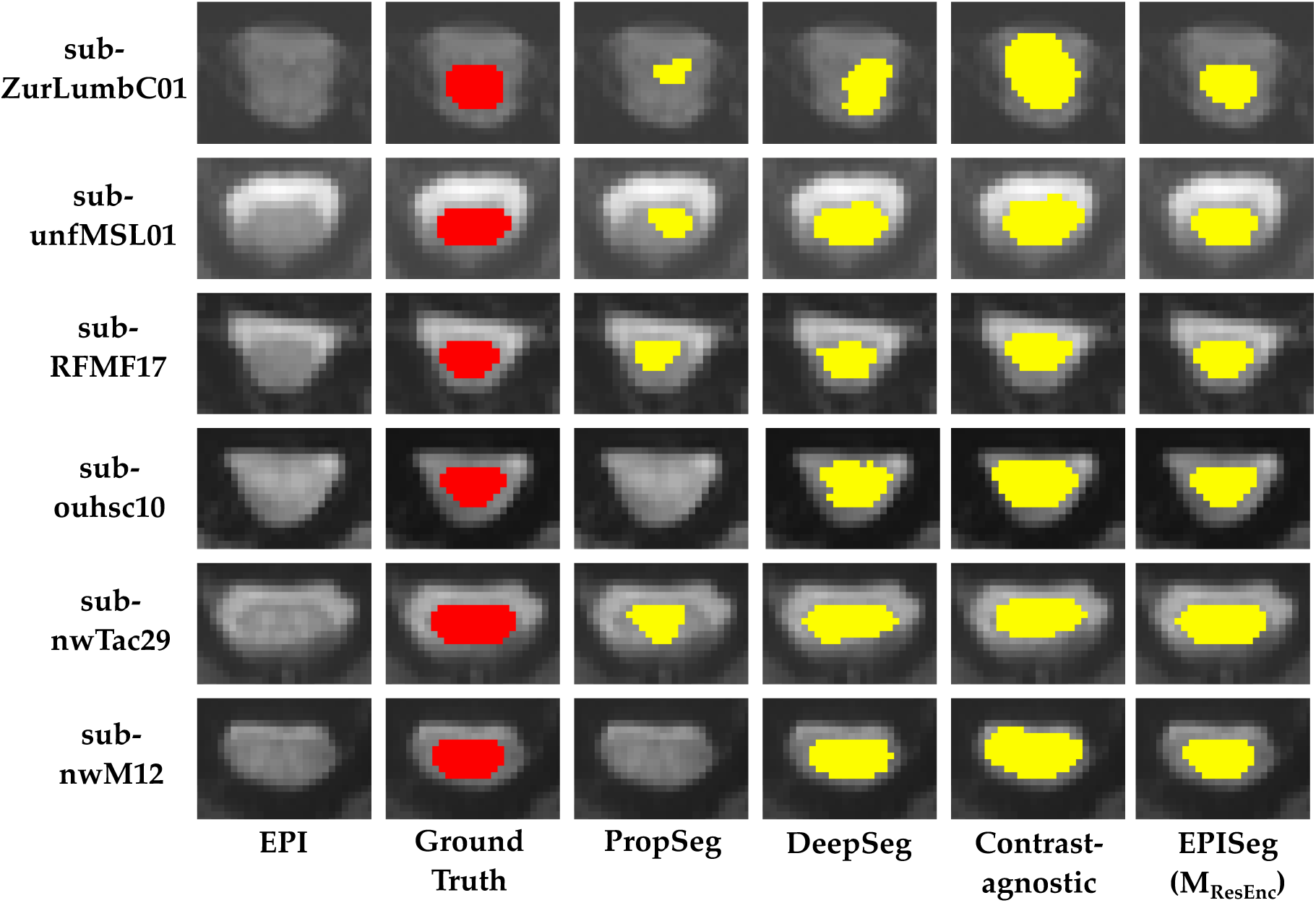
Visual assessment. Comparison of EPISeg (*M_ResEnc_*) with other automatic SC segmentation methods. The colored overlay shows the SC on the images. We can observe that EPISeg provides the most precise segmentations.

### 3.4 Qualitative assessment

We evaluated our model’s performance on *D_test_* through a qualitative assessment involving two raters with 6 and 18 years of experience of SC MRI experience, respectively. These raters meticulously examined the segmentation results for each slice, as documented in the QC report generated by the sct qc functionality in SCT. Figure 5 showcases the performance of various models on selected axial slices, highlighting the quantitative performance of each. The participants represented in the figure include healthy controls as well as patients with degenerative cervical myelopathy and fibromyalgia.

In our evaluations, we observed that PropSeg frequently under-segments and sometimes fails to segment the SC entirely, especially when the SC is not clearly distinguishable from the cerebrospinal fluid. DeepSeg performs slightly better but still displays inconsistencies in SC segmentation. The Contrast-agnostic model offers more consistent segmentation boundaries but often over-segments the SC. EPISeg demonstrates superior performance, closely matching the ground truth. For instance, EPISeg successfully segmented the SC in the sub-ZurLumbC01, a dataset with low-quality lumbar cord images that were not included in the training set.

## 4 Discussion

In this study, we introduce EPISeg, a DL-based method for automatic SC segmentation from gradient-echo EPI images, which are widely used in (fMRI). EPISeg is robust to heterogeneous resolutions, scanner settings, and motion artifacts. Consequently, it addresses challenges that have historically hindered SC segmentation on EPI data, requiring labor-intensive and bias-prone manual interventions.

In order to develop a robust EPI segmentation model, the first necessary step was to create a dataset that would provide a DL model with enough context to be able to segment the SC in a low-quality image setting. We curated a large multi-site EPI SC dataset consisting of spinal and corticospinal acquisitions, arranged it as per the BIDS convention, and made it publicly available in OpenNeuro - https://openneuro.org/datasets/ds005143/versions/1.3.0.

While creating the dataset, we observed that the provided ground truth was erroneous in some cases (e.g., over-segmented or under-segmented). This led us to the decision to train EPISeg sequentially. We used good-quality segmentations to train a strong baseline model which provided high-quality segmentations before successive models. At each model training step, a pool of best predictions was chosen along with manual correction wherever required, including verification by an expert. We obtained our final model using these steps iteratively, which was able to segment the SC reliably. We demonstrated the robustness of our model against low spatial resolution of EPIs, susceptibility artifacts, signal drop-outs, ghosting, and motion-related distortions and eliminated the need for manual interventions.

In terms of SC segmentation directly on EPI data, only [Giulietti et al., 2011] provides a semi-automatic method. This method consists of multiple preprocessing steps before the SC was segmented using k-means clustering. The authors claim that their method’s performance efficiency declines with the increase in the distortion present in the image. Even though we cannot reproduce this method for comparison due to the lack of provided details regarding the parameters used, we show that, in addition to being completely automatic, EPIseg is highly robust to distortions present in EPI data.

Our method, EPISeg, demonstrates superior accuracy and robustness in spinal cord segmentation compared to state-of-the-art models like PropSeg, DeepSeg, and Contrast-agnostic segmentation. It generalizes well across different SC conditions and handles out-of-distribution data effectively. EPISeg achieves a mean Dice score of 0.88 with a standard deviation of 0.05, reflecting its consistent performance. PropSeg performs poorly as it was primarily optimized for T1 and T2 contrasts and is not well-suited for EPI data. Similarly, DeepSeg and the contrast-agnostic model, which were not specifically trained for spinal cord segmentation on EPI images, lack sufficient context to accurately handle the unique characteristics of EPI data. Although the Contrast-agnostic model performs better than PropSeg and DeepSeg due to its broader training dataset, it still falls short of EPISeg’s performance.

To potentially circumvent the need for segmenting the EPI for registration purposes, a high-resolution anatomical image can be acquired with the same field-of-view as the EPI images (in a way that registration between this image and the EPI is not necessary). However, this approach has also drawbacks. Mainly, the acquisition of a high-resolution image would require additional scan time. Especially considering the movement of a participant within and across multiple functional runs, it may be even necessary to acquire one high-resolution image which would significantly increase scan time.

Despite the superior performance of EPISeg for segmentation of spinal cord fMRI data against other automated methods, several limitations of the current work must be considered. Although this study includes a wide range of data from multiple sites and participants including data from different patient populations, it does not cover all possible data domains. Variations in acquisition parameters and spinal cord conditions across different populations may not be fully represented here, possibly limiting the generalizability of our findings. Additionally, our method has solely been tested on data acquired at 3T. Segmentation performance and accuracy could vary across other field strengths, due to the fact that, at lower (1.5T) and higher (7T) fields, differences in signal-to-noise ratio, contrast, and spatial resolution may influence the quality of the EPI data.

EPISeg here is optimized for EPI sequences as they are the most commonly used technique for fMRI acquisitions. However, spinal cord fMRI can also be performed using other sequence types (such as T2*-weighted fast gradient echo acquisitions). The performance of EPIseg on other acquisition types has not been evaluated in this study. Finally, although we gathered data from various sites that have employed different motion correction techniques, it is important to remember that effective motion correction is critical to the success of spinal cord fMRI segmentation, given the susceptibility of this region to motion artifacts from respiration, pulsation, and participant movement. Inadequate motion correction could impact the accuracy of segmentation by obscuring spinal cord boundaries. Future work should aim to broaden the data domains represented, including a wider range of acquisition parameters, populations, and magnetic field strengths (1.5T, 3T, and 7T). Moreover, validation of EPIseg on GRE T2*-weighted sequences and improvements in motion correction strategies will be necessary to enhance segmentation robustness and ensure that it performs effectively across diverse imaging conditions.

## 5 Conclusion

We provide an automatic SC segmentation method for gradient-echo EPI. We demonstrate the performance of our method on various spinal and also corticospinal fMRI datasets (including samples from clinical populations) from different sites with different acquisition parameters. The dataset is publicly available on OpenNeuro. This reliable segmentation method will provide a strong basis for fMRI data preprocessing, particularly for spatial normalization which is necessary for obtaining reliable group-level results. Our method is readily available as a part of SCT v6.3 or higher.

## Data and Code Availability

All the data used for this study is open-sourced and can be found at: https://openneuro.org/datasets/ds005143/versions/1.3.0

All the scripts, instructions to reproduce results and the lastest model weights can be found here: https://github.com/sct-pipeline/fmri-segmentation/releases/tag/v0.2. This method is integrated into SCT v6.3 and can be used by using the *sct deepseg* command with the *-task seg sc epi* parameter.

## Author Contributions

Rohan Banerjee: Data Curation, Methodology, Formal Analysis, Software, Validation, Writing – Original Draft, Writing – Review Editing, Visualization; Merve Kaptan: Data Curation, Validation, Methodology, Writing – Original Draft, Writing – Review Editing; Kenneth A. Weber II: Supervision, Writing – Review Editing, Conceptualization, Funding Acquisition; Benjamin De Leener: Supervision, Methodology, Writing – Review Editing, Conceptualization, Funding Acquisition; Julien Cohen-Adad: Supervision, Methodology, Writing – Review Editing, Conceptualization, Funding Acquisition; Alexandra Tinnermann: Data Contribution; Ali Khatibi: Data Contribution; Alice Dabbagh: Data Contribution; Christian Büchel: Data Contribution; Christian W. Kündig: Data Contribution; Christine S.W. Law: Data Contribution; Dario Pfyffer: Data Contribution, Writing – Review Editing; David J. Lythgoe: Data Contribution, Writing – Review Editing; Dimitra Tsivaka: Data Contribution; Dimitri Van De Ville: Data Contribution; Falk Eippert: Data Contribution, Writing – Review Editing; Fauziyya Muhammad: Data Contribution; Gary H. Glover: Funding Acquisition, Data Contribution; Gergely David: Data Contribution, Writing – Review Editing; Grace Haynes: Data Contribution; Jan Haaker: Data Contribution, Writing – Review Editing; Jonathan C. W. Brooks: Data Contribution, Writing – Review Editing; Jürgen Finsterbusch: Data Contribution, Writing – Review Editing; Katherine T. Martucci: Data Contribution, Funding Acquisition, Writing – Review Editing; Kimberly J. Hemmerling: Data Contribution, Mahdi Mobarak-Abadi: Data Contribution; Mark A. Hoggarth:Data Contribution; Matthew A. Howard: Data Contribution; Molly G. Bright: Data Contribution; Nawal Kinany: Data Contribution; Olivia S. Kowalczyk: Data Contribution; Patrick Freund: Data Contribution; Robert L. Barry: Data Contribution, Funding Acquisition, Writing – Review Editing; Sean Mackey: Funding Acquisition, Data Contribution; Simon Schading: Data Contribution; Shahabeddin Vahdat: Data Contribution; Todd Parish: Data Contribution; Yufen Chen: Data Contribution; Véronique Marchand-Pauvert: Data Contribution; Zachary A. Smith: Data Contribution

## Acknowledgements

We would like to thank Jan Valosek for their valuable feedback and review of the manuscript. We would also like to thank Joshua Newton and Mathieu Guay-Paquet for helping us integrate our EPISeg to SCT. We express our gratitude towards the members of the NeuroPoly Lab at Polytechnique Montreal for all the fruitful discussions.

